# Whole genome-based characterization of multi-drug resistant *Enterobacter* and *Klebsiella aerogenes* isolates from Lebanon

**DOI:** 10.1101/2022.06.17.496657

**Authors:** Georgi Merhi, Sara Amayri, Ibrahim Bitar, George F. Araj, Sima Tokajian

## Abstract

**Background:** Enterobacter spp. are rod-shaped Gram-negative opportunistic pathogens belonging to Enterobacterales. This study aimed at the molecular and genomic characterization of multi-drug resistant *Enterobacter* spp. isolates recovered from hospitalized patients in a tertiary care hospital in Lebanon.

**Materials:** A total of 59 *Enterobacter* spp. clinical isolates consisting of 41 carbapenem-resistant and 18 susceptible by E-test were included in this study. Genotypic identification through whole-genome sequencing was performed and confirmed *in silico*. Resistance and plasmid profiles were studied using ResFinder4.0 and Plasmid-Finder2.1. Multi-locus sequence typing (MLST) was used to determine the isolates’ clonality.

**Results:** ANI identified and confirmed that 47 (80%) isolates were *E. hormaechei*, 11 (18%) were *Klebsiella aerogenes* and 1 (2%) was an *E. cloacae*. Carbapenem-resistance was detected among 41 isolates all showing an MIC_90_ of ≥ 32 µg/ml for ertapenem, imipenem, and meropenem. *bla*_NDM-1_ (58.5%), *bla*_*ACT*_*-*_16_ (54%), and *bla*_OXA-1_ (54%) were the most common detected β-lactamases, while *bla*_CTX-M-15_ gene (68%) was the main detected extended-spectrum β-lactamase (ESBL) encoding gene. Chromosomal *ampC* gene, carbapenemase encoding genes, and porin modifications were among the detected carbapenem resistance determinants. The carbapenemase encoding genes were linked to three well-defined plasmid Inc groups, IncFII/IncFIB, IncX3, and IncL. MLST typing revealed the diversity within the studied isolates, with ST114 being the most common amongst the studied *E. hormaechei*.

**Conclusion:** The spread of carbapenem-resistant isolates in clinical settings in Lebanon is a serious challenge. Screening and continuous monitoring through WGS analysis could effectively limit the dissemination of drug-resistant isolates in hospitalized patients.

**Importance:** Drug resistance is an increasing global public health threat that involves most disease-causing organisms and antimicrobial drugs. Drug-resistant organisms spread in healthcare settings, and resistance to multiple drugs is common. Our study demonstrated the mechanisms leading to resistance against the last resort antimicrobial agents among members of the *Enterobacteriaceae* family. The spread of carbapenem-resistant bacteria in clinical settings is a serious challenge. Screening and continuous monitoring could effectively limit the dissemination of drug-resistant isolates in hospitalized patients.

## Introduction

*Enterobacter cloacae* complex (ECC) members and *Klebsiella aerogenes*, belong to the family *Enterobacteriaceae* and are Gram-negative, rod-shaped, non-spore forming opportunistic pathogens causing healthcare-associated infections (HAIs) (1–3). ECC members are saprophytic microorganisms in that they inhabit and colonize diverse environments such as sewage, soil and the human GI tract (4). Detection of *K. aerogenes* is mostly linked to clinical settings and geographically widespread outbreaks (2). The emergence of ECC and *K. aerogenes* as potent nosocomial pathogens was further aggravated by the worldwide occurrence of multi-drug resistant clones (5).

Hoffmann *et al*, separated the ECC into 12 (I to XII) diverse genetic clusters based on *hsp60* single marker sequencing (6). Currently, *hsp60* typing remains the most widely used method categorizing species within the ECC (5). With the advent of whole-genome sequencing however, average nucleotide identity (ANI) analysis is becoming the gold standard for *in silico* species assignment. Using such approaches on the entire collection of RefSeq *Enterobacter* genomes allowed the distinction of 18 phylogenetic clusters (A to R) within the ECC (7).

Carbapenems are considered drugs of last resort for the treatment of extended-spectrum β-lactamase (ESBL) producing Enterobacterales isolates (8). ECC members were among the first Enterobacterales to harbor carbapenem resistance determinants and are currently the second most prevalent Carbapenem resistant *Enterobacteriaceae* (CRE) in the US and other major regions (3). Carbapenemase producing *Enterobacter* spp. (CPE) isolates in hospital acquired infections (HAIs) predominately belonged to the *E. hormaechei* species with its various respective subspecies (3, 9). *E. hormaechei* subsp. *xiangfangensis* (CIII) and *E. hormaechei* subsp. *steigerwaltii* (ST114, ST90 and ST93) are the most geographically prevalent CPEs (9).

There is a wide data schism regarding the epidemiology and dissemination of clinically significant *K. aerogenes* and It’s only lately that interest in this pathogen is rising. However, data related to the global burden of resistant subtypes of this pathogen are still scarce, especially in the Middle East (10). Well documented clinical cases of carbapenem resistant *K. aerogenes* (CR-KE) in the US, Europe and Asia were linked to plasmid encoding carbapenemases (2). Carbapenem resistance mechanism in CR-KE were mainly linked to the constitutive overexpression of chromosomal AmpC along with changes in membrane permeability through porin regulation (10).

The ominous antimicrobial resistance (AMR) phenomenon has become a global issue in general and Lebanon in particular (11). Various studies in Lebanon described alarming cases of carbapenem resistance in members of the family *Enterobacteriaceae*. Arabaghian *et al*, reported a 70.6% rate of total carbapenem resistance in a *Klebsiella pneumoniae* isolate set from a tertiary care hospital (12), and Alousi *et al*, reported the first ST-405 *Escherichia coli* single isolate carrying the *bla*_OXA-48_ carbapenem resistance determinant (13). Despite their notoriety as highly resistant nosocomial pathogens, members of *Enterobacter* spp. were not well studied. The sporadic studies that tackled their dissemination and resistance patterns, were based on phenotypic screening and PCR assays (14, 15). A recently published comprehensive statistical report about ESBL and carbapenemase carriage in Lebanon for ESKAPE group pathogens also lacked detailed molecular characterization (16).

We aimed in this study to setup a workflow for the accurate processing of ECC and *K. aerogenes* using isolates recovered from Lebanon. To that end, we performed a detailed molecular characterization of 50 multi-drug resistant (MDR) *E. cloacae* complex and *K. aerogenes* clinical isolates. We relied on whole-genome sequencing (WGS) to elucidate their correct nomenclature, clonal relatedness, and resistance/virulence profiles. The sequenced genomes were subjected to both intra- and inter-analysis which allowed us to build an in-depth genomic understanding of ECC and *K. aerogenes* isolates recovered from Lebanon and in positioning them within a larger more comprehensive global context. This study also included detailed analysis pertaining to the pathogenesis and dissemination of the re-emerging pathogen *K. aerogenes* in Lebanon.

## Materials and Methods

### Ethical approval

Ethical approval for this project was not required as the study isolates were collected and stored as part of routine clinical screening. Patients’ records and information were anonymized, and clinical isolates data were de-identified before undertaking any form of analysis.

### Bacterial Isolates

A total of 59 ECC and *K. aerogenes* isolates were collected from the 350-bed tertiary care teaching hospital AUBMC as part of routine clinical care screening. Preliminary species identification was performed through the Matrix Assisted Laser Desorption/Ionization Time of Flight (MALDI-TOF) system (Bruker Daltonik, GmbH, Bremen, Germany) following the manufacturer’s instructions. 81% (48/59) were identified as members of the EC complex whereas 19% (11/59) belonged to the *K. aerogenes* species and accordingly were designated as ENM1-48 and KAM1-11, respectively. Isolates were recovered from different infection sites sources: skin, urine, wounds and abdominal fluids. The mean patients’ age was 47 ± 26 years old with the range being 20 days to 90 years old.

### Antibiotic susceptibility testing

Antibiotic susceptibility testing was performed through the Kirby-Bauer disk diffusion test (DDT) on Muller-Hinton agar including an assortment of 18 antibiotic disks representing 8 categories of regularly used antimicrobial agents in clinical settings. The minimum inhibitory concentration (MIC) was determined for 8 selected antibiotics. All results were interpreted following CLSI recommendations (17, 18).

### Bacterial DNA extraction

Colonies were grown overnight on TSA agar plates. The Nucleospin® Tissue kit (Macherey-Nagel, Germany) was used for bacterial DNA extraction according to the manufacturer’s instructions. The obtained DNA was used for downstream PCR assays and whole-genome sequencing.

### *Hsp60* gene amplification and sequencing

PCR amplification and sequencing were performed with the *hsp60*-F and *hsp60*-R primers as previously described (6). *hsp60* sequences were aligned with references representing the 12 Hoffmann clusters (I→ XII) using Clustal-Omega alignment tool, (https://www.ebi.ac.uk/Tools/msa/clustalo/) (19). The Hoffman cluster neighbor-joining (NJ) tree was visualized using the interactive tree of life (iTOL) tool (20).

### Pulse Field Gel Electrophoresis (PFGE)

PFGE was carried out using *Xba*I restriction enzyme (ThermoScientific, Waltham, MA, USA), according to the pulseNet standard protocol (http://www.pulsenetinternational.org; https://www.cdc.gov/pulsenet/). Fingerprint analysis was performed using BioNumerics software version 7.6.1 (Applied Maths, Sint-Martens-Latem, Belgium) and PFGE profiles were categorized as pulsotypes based on the criteria of a minimal difference of three or more bands between select isolates (21). Fingerprint clusters were inferred with the BioNumerics software based on the Dice correlation with optimization and tolerance values set at 1.5%.

### Multi-Locus sequence typing (MLST)

MLST was carried out for ECC members as previously described by Miyoshi-Akiyama *et al*. (22) and resulting sequences were curated versus the public database for *E. cloacae* complex isolates (https://pubmlst.org/ecloacae/). *K. aerogenes* isolates sequence types (ST) were also determined using the publicly available database for the molecular typing of *K. aerogenes* (https://pubmlst.org/kaerogenes/).

### Plasmid Based Replicon Typing (PBRT)

Plasmid incompatibility (Inc) types were elucidated using the DIATHEVA PBRT kit (Diatheva, Fano, Italy) through a PCR-based replicon typing method consisting of eight multiplex PCR assays for the amplification of 25 replicons. All reactions were carried out according to the manufacturer’s instructions, including positive controls for all assays.

### Conjugative Transfer

The transfer of resistant markers (*bla*_NDM-1_, *bla*_NDM-5_, *bla*_OXA-48_, and *bla*_OXA-181_) was determined by conjugation using a plasmid-free recipient azide resistant strain *E. coli* J53 and the study isolates as donors. Single colonies of both the donor and recipient were inoculated on Luria-bertani (LB) broth and incubated overnight at 37°C. Equal volumes of the donor and the recipient cultures were mixed and incubated overnight at 37°C, followed by plating on LB agar supplemented with meropenem. To ensure the effectiveness of the selected plates or the used antibiotics, positive and negative controls were run in parallel by separately plating the donor and recipient. Presence or absence of carbapenemases was determined through a PCR assay on the trans-conjugants followed by PCR-Based Replicon Typing (PBRT).

### Whole-Genome sequencing and Quality Control (QC)

Genomic libraries of all 59 study isolates were created using the Nextera XT DNA library preparation kit with dual indexing (Illumina). The resulting libraries were sequenced on an Illumina MiSeq with forward and reverse reads with 300 bp length. Long read sequencing technology was applied on the PacBio Sequel I platform (Pacific Biosciences, Menlo Park, CA, USA) for three representative isolates ENM17, ENM30 and KAM9 as previously described (23). Quality assessment for the raw reads was done using FastQC version 0.11.8 and QUAST version 5.0.2 (24, 25). Adapter removal and trimming was performed through Trimmomatic v0.39 (26).

### Assembly, identification, annotation and genome analysis

Long read sequencing data were assembled using the HGAP4 De Novo Assembly Application (Pacific Biosciences) with a minimum seed coverage of 30 X. Short read trimmed sequences were assembled with SPAdes version 3.14 with read error correction enabled (27).

Accurate species identification was performed through Average Nucleotide Identity (ANI) testing. FastANI tool was used to generate a distance matrix of ECC isolates against Refseq complete *E. hormaechei* and *E. cloacae* genomes (supplementary table S2) (28). Subsequently, the assembled genomes were uploaded and annotated with the RASTtk (http://rast.nmpdr.org). ResFinder 4.1 (https://cge.cbs.dtu.dk/services/ResFinder/) and the comprehensive antibiotic resistance database (CARD) were used to assess resistance determinants in all the isolates (29, 30). Sequences of the *omp35* and *omp36* porin genes were extracted from the *K. aerogenes* genomes and sequences were translated to detect non-synonymous mutations, using the translate webtool available at the Bioinformatics resource portal (https://web.expasy.org/translate/). Putative K-types and virulence systems (yersiniabactin and colibactin) were inferred with the Kleborate and Kaptive (31).

### Plasmid studies

PlasmidFinder 2.1 was used to detect and confirm the plasmid Inc types found in the study isolates (32). Potential plasmidic contigs resulting from long read sequencing were extracted and the closest matching reference was detected through the BLASTn tool on NCBI (www.ncbi.nlm.nih.gov/BLAST). PLACNETw was also used to identify plasmid sequences from short read sequencing data (33). Sequences were aligned and overlapping bases were trimmed and contiguous sequences were manually assembled. Open Reading Frames (ORFs) were inferred with Prodigal v2.6.1 with the closed option enabled (34). All coding sequences were investigated and manually annotated through a BLASTp strategy and the IS finder database for insertion sequences (https://isfinder.biotoul.fr/).

### Phylogenetic Analysis

The GToTree pipeline was used to determine the phylogenetic relationship between the study isolates (ECC and *K. aerogenes*). The pipeline relies on the analysis of 172 single-copy genes (SCGs) unique to Gamma-proteobacteria and outputs a phylogenetic tree based on the comparison of the selected SCG set within the specified input files (35). All ECC and *K. aerogenes* complete Refseq genomes were downloaded from NCBI with the “complete genome” and “latest Refseq” filters. The final tree was generated and visualized on iTOL (20).

### Sequence Data

All 59 WGS sequences were deposited at DDBJ/EMBL/GenBank under the BioProject number **PRJNA551102**. Plasmid sequences of pLAU_ENM30_NDM1, pLAU_ENM17_OXA181 and pLAU_KAM9_OXA48 were deposited at DDBJ/EMBL/GenBank under the following accession numbers: **MN792917, MN792918** and **MN792919**.

## Results

### Characterization of the Study Isolates

Isolates were collected over a period of five years (2013-2018) as part of routine microbiological screening at the American University of Beirut Medical Center (AUBMC). Primary criterion for selection was based on non-susceptibility (intermediate/resistant) to one or more of the three clinically used carbapenems (meropenem, ertapenem and imipenem). Secondary inclusion criterion was any isolate recovered from an infection displaying a susceptible phenotype to the clinically tested antibiotics. Most of the isolates (72 %; n=43/59) displayed resistance patterns to at least two and more out of the three tested carbapenems and 9% (4/43) of sub-selected isolates showed non-susceptibility to two tested carbapenems. Only 7% (3/43) were resistant to only one. All non-susceptible isolates (n=43) were resistant to ertapenem out of which four additionally were resistant to meropenem (KAM8/9-ENM28/29). KAM1, KAM4 and ENM48 were resistant only to ertapenem. Additionally, more than 60% of the isolates showed resistance to other antibiotic classes; 75% (44/59) were resistant to gentamycin and 61% (36/59) to Trimethoprim-sulfamethoxazole.

ResFinder, used for the *in silico* analysis of antibiotic resistant determinants (ARD), uncovered the presence of 60 different unique resistance genes. The detected genes represented the nine clinically used antibacterial classes and which included: β-lactams, aminoglycosides, fluroquinolones, trimethoprim, sulfonamide, Fosfomycin, tetracyclines, macrolides and chloramphenicol (Figure 1). Chromosomal *ampC* genes were detected in all the ECC isolates.

**Figure 1.**
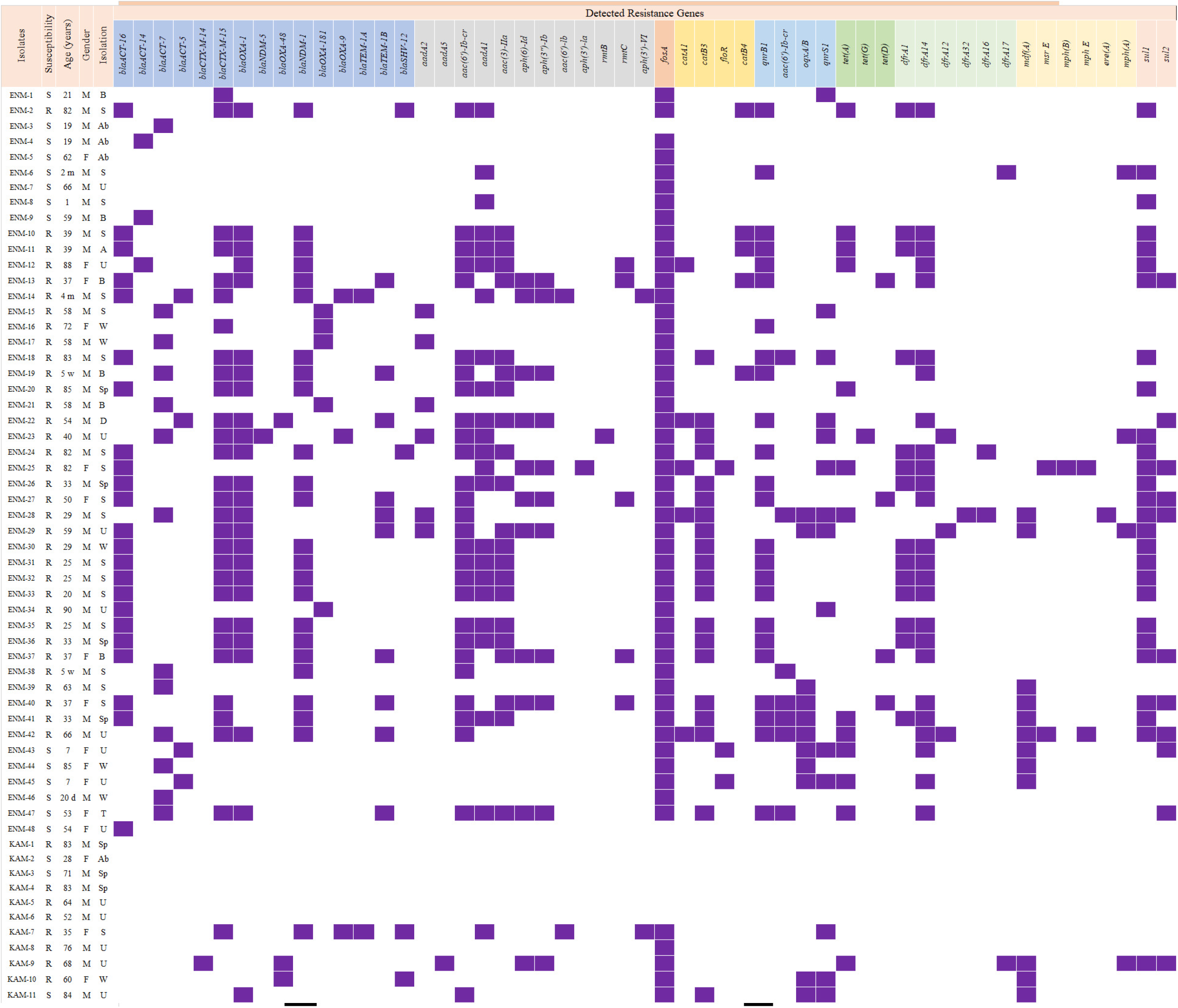
Isolates’ information and detected antibiotic resistance genes. In the susceptibility column, R: Resistant to Carbapenem; S: susceptible to Carbapenem. In the Age column, d: days m: months; w: weeks. In the gender column, F: Female; M: Male. In the specimen isolation column, A: Abscess; Ab: Abdominal fluid; B: Blood; D: DTA; S: Skin; Sp: Sputum; U: Urine; W: Wound.

The prevalence of carbapenem resistance determinants (CRDs), shown in Figure 1, was validated using *in silico* whole-genome sequence analysis. *bla*_NDM-1_ was dominant and detected in 41% (24/59) of the isolates. Second-most dominant gene was the class D oxacillinase *bla*_OXA-181_ at 8% (5/59) followed closely by *bla*_OXA-48_ at 5% (3/59). *bla*_NDM-5_ was only detected in one ECC isolate (2%; ENM23). *bla*_ACT-16_ and *bla*_ACT-7_ were the most common *ampC* genes at 39% (23/59) and 22% (13/59), respectively. Amongst the genes conferring the ESBL phenotype, *bla*_CTX-M-15_ and *bla*_OXA-1_ were the most observed variants in this study, with *bla*_CTX-M-15_ being present in 49% (29/59) of isolates and *bla*_OXA-1_ in 41% (24/59), and their presence was not mutually exclusive. Seven isolates (KAM5, KAM6, KAM8, ENM25, ENM28/29 and ENM39) did not harbor any carbapenemase encoding gene. ENM-25 and ENM-39 only harbored their respective chromosomal *ampC* determinant suggesting a revertant wild type phenotype. In contrast, both ENM-28 and ENM-29 harbored ESBL encoding genes (Figure 1).

Analysis of outer membrane protein encoding genes against functional references (CP017179.1, CP017179.1, CP017179.1, AF335467.1) showed that only KAM-5 and KAM-6 had a mutated form of the *omp36* gene with a premature insertion of an Amber stop codon. The remaining six isolates however, had functional *omp* genes with the absence of any potential non-synonymous mutation.

All three *K. aerogenes* isolates were devoid of any β-lactamases encoding genes.

### *Hsp60* Typing and Average nucleotide identity analysis

MALDI-TOF identification placed 81% (48/59) of the isolates under the *E. cloacae* complex. Subsequently, ECC isolates were subjected to *hsp60* typing. The *hsp60* sequences of the 48 ECC members were aligned with type strain sequences, from the NCBI gene database (AJ417141-3, AJ543870, AJ543878, AJ543807, AJ417110, AJ546761, AJ417134, AJ567879, AJ417115 and AJ862866). 15% (7/48) of ECC isolates matched with the *E. cloacae* cluster III, whereas only one isolate matched to the *E. cloacae* subsp. *cloacae* mini clade (Figure 2). It is noteworthy that 84% (40/48) of the isolates clustered with *E. hormaechei* type strain sequences (AJ567884.1, AJ567885.1, AJ567878.1), whereas none was typed as subsp. *hormaechei*. The remaining 40 were distributed between the *steigerwaltii* (12/48, 25%) or the *oharae* subspecies (28/48, 58%) (Figure 2).

**Figure 2.**
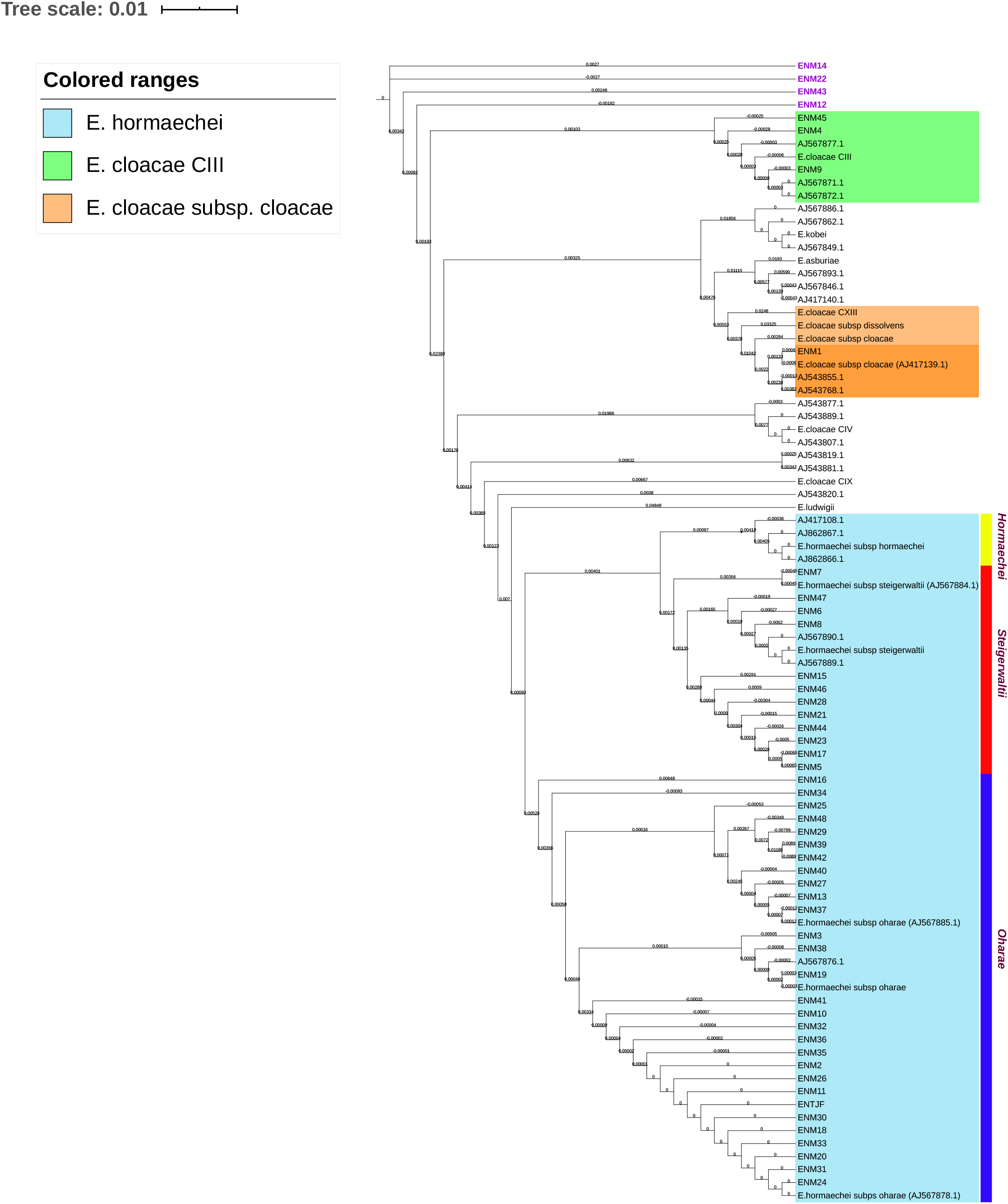
Neighbor-Joining (NJ) tree based on multisequence alignment of partial *hsp60* gene for all ECC isolates. Isolates clustering with reference *hsp60* genes were highlight within colored ranges. *Hormaechei* subspecies were demarcated through colored strips. Isolates ENM12-14-22-43 were highlighted in purple as each has clustered as a singleton on the NJ tree.

Average nucleotide identity (ANI) testing was employed as a confirmatory and higher sensitivity test for species and sub-species identification. Eight refseq complete genomes (Supplementary Table S1) were used as references representing *K. aerogenes, E. cloacae* spp. *cloacae*/*dissolvens* and the five subspecies of *E. hormaechei* (*xiangfangensis, hoffmannii, oharae, steigerwaltii, hormaechei*). ANI values shown in Supplementary Table S2, confirmed the identification of the 11 *K. aerogenes* isolates. As for members of the ECC, results obtained revealed significant discrepancies between *hsp60* typing and ANI testing. According to the results of this study, the predominant *hormaechei* subsp. was *xiangfangensis* at 54% (26/48) followed by 27% (13/48) and 15% (7/48) for *steigerwaltii* and *hoffmannii*, respectively. Only one *E. hormaechei* isolate (ENM-48) belonged to the subsp. *oharae* and, similarly, only one isolate was *E. cloacae* subsp. *cloaca* (ENM-1).

### Typing

Twenty-four different sequence types (STs) were detected among the EC complex. Fourteen of the isolates were of ST114 (29.2%), five of ST182 (10.4%), and three of ST190 (6.25%). With *K. aerogenes*, seven distinct sequence types (STs) were detected including: ST143 (18.2%; 2/11), ST93 (18.2%; 2/11), ST209 (18.2%; 2/11) and ST210 (18.2%; 2/11) (Figure 3).

**Figure 3.**
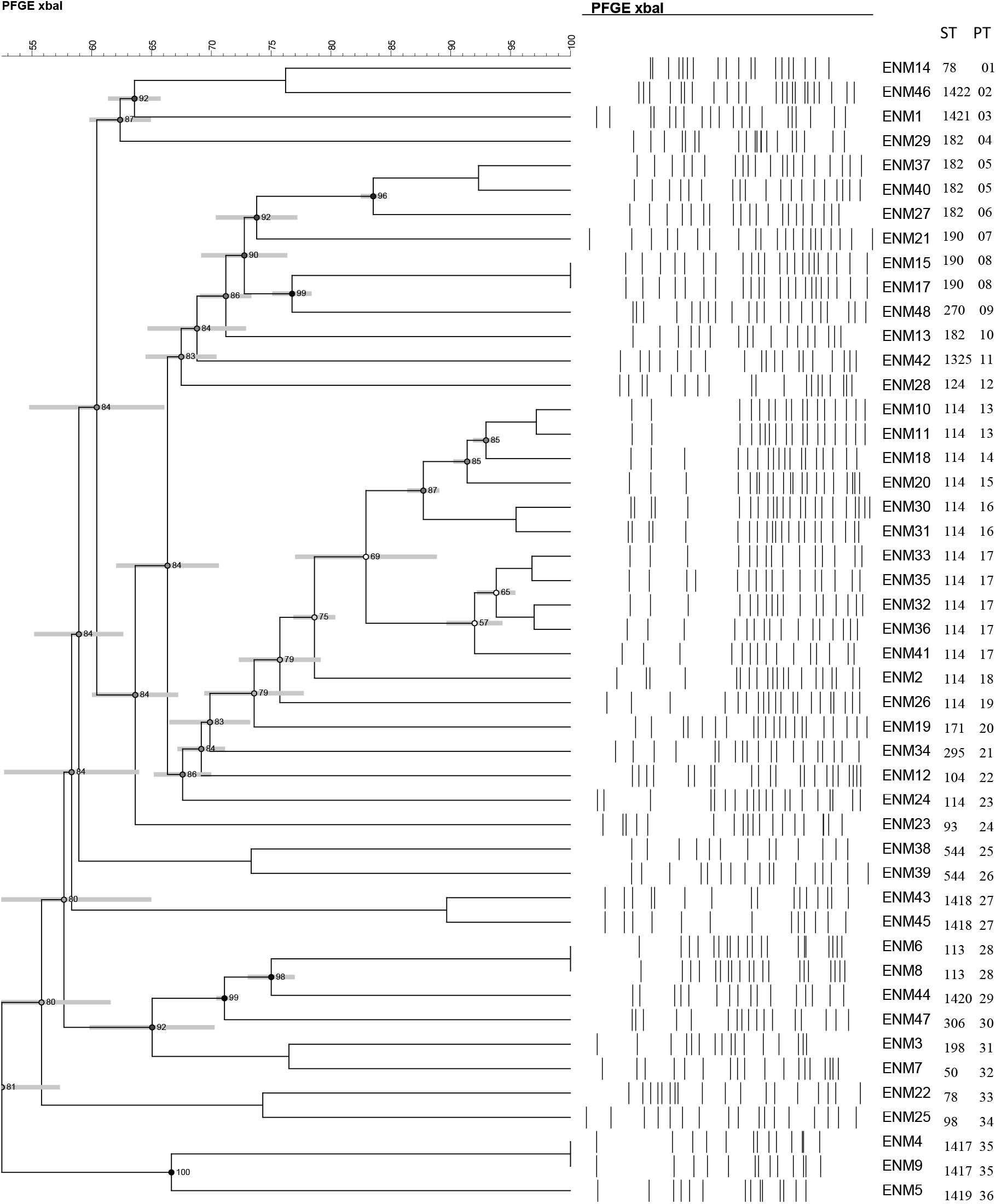
PFGE dendrogram of ECC isolates. Dendrogram was generated by BioNumerics software version 7.6.1 showing isolate clonality based on their banding patterns generated by XbaI restriction digestion. The results were inferred using the dice correlation with optimization and tolerance values set at 1.5% Isolates’ STs, and pulsotypes (PT) are, also, shown.

Pulse-field Gel electrophoresis was performed to study the genetic relatedness between the isolates. A total of 36 distinct pulsotypes were identified. ENM-37 and ENM-40 showed a similarity of 93% based on the banding patterns (Figure 3). These two isolates are of the same subsp. (*E. hormaechei* subsp. *xiangfangensis*) belonging to the sequence type ST182, and with mostly identical resistance and plasmid profiles. ST114 isolates however, had seven different pulsotypes where the PFGE banding patterns did not match fully with the MLST typing. All ST114 isolates shared ≥80% PFGE similarity patterns with identical resistance patterns and determinants, except for ENM-24 which had a distinct PFGE pattern compared to all other ST114 isolates. ENM-16 was untypable even after treatment with secondary enzyme.

Seven different pulsotypes were determined based on clustering and banding patterns for *K. aerogenes* (Figure 4). Some of the banding patterns matched with the MLST. KAM-9 and KAM-10 are of ST143 and shared high degree of clonal relatedness with only one band difference. KAM-2 and KAM-11 on the other hand, were typed as ST93 and distinct banding patterns.

**Figure 4.**
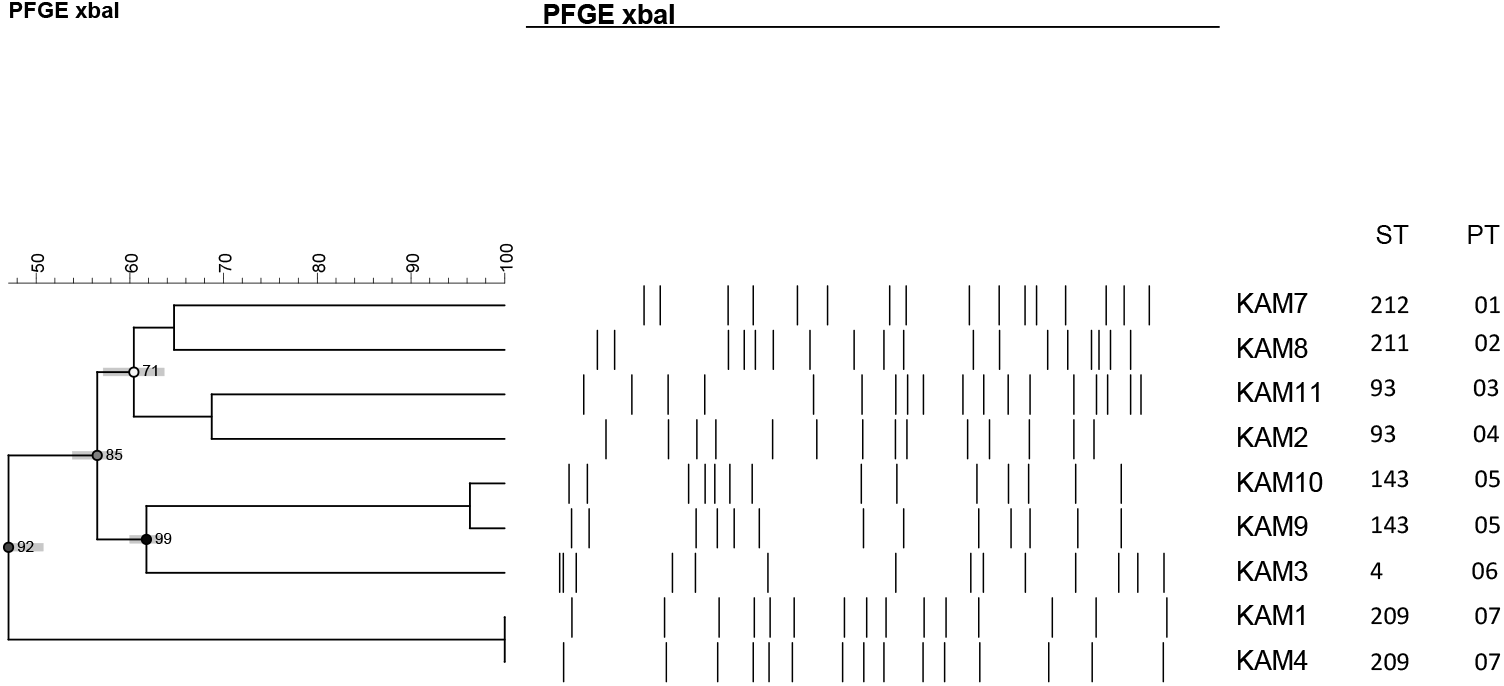
PFGE dendrogram of *K. aerogenes* isolates. Dendrogram was generated by BioNumerics software version 7.6.1 showing isolate clonality based on their banding patterns generated by XbaI restriction digestion. The results were inferred using the dice correlation with optimization and tolerance values set at 1.5% Isolates’ STs, and pulsotypes (PT) are, also, shown. Isolates KAM5 and KAM6 were untypable.

We used Kaptive, a web-based tool for the *in silico* k-type determination through targeting the complete *cps* locus, and accordingly we detected three different K-types among the eleven *K. aerogenes* isolates. All the *K. aerogenes* were identified as having the KL68 K-type except for KAM-3 and KAM-8, which respectively were identified as KL107 and KL26 (Table S3).

### Phylogenetic Analysis

The primary phylogenetic placement of the isolates was performed through the GToTree pipeline. The analysis of 172 SCGs for all the isolates and complete RefSeq genomes clustered all into two major clades (I & II) (Figure 5). Clade I (red block) distinctively harbored all the *K. aerogenes* (11/59) and at a large distance from the remaining *Enterobacter* spp. isolates (Figure 5). All the ENM isolates (ENM1-48) clustered within clade II.

**Figure 5.**
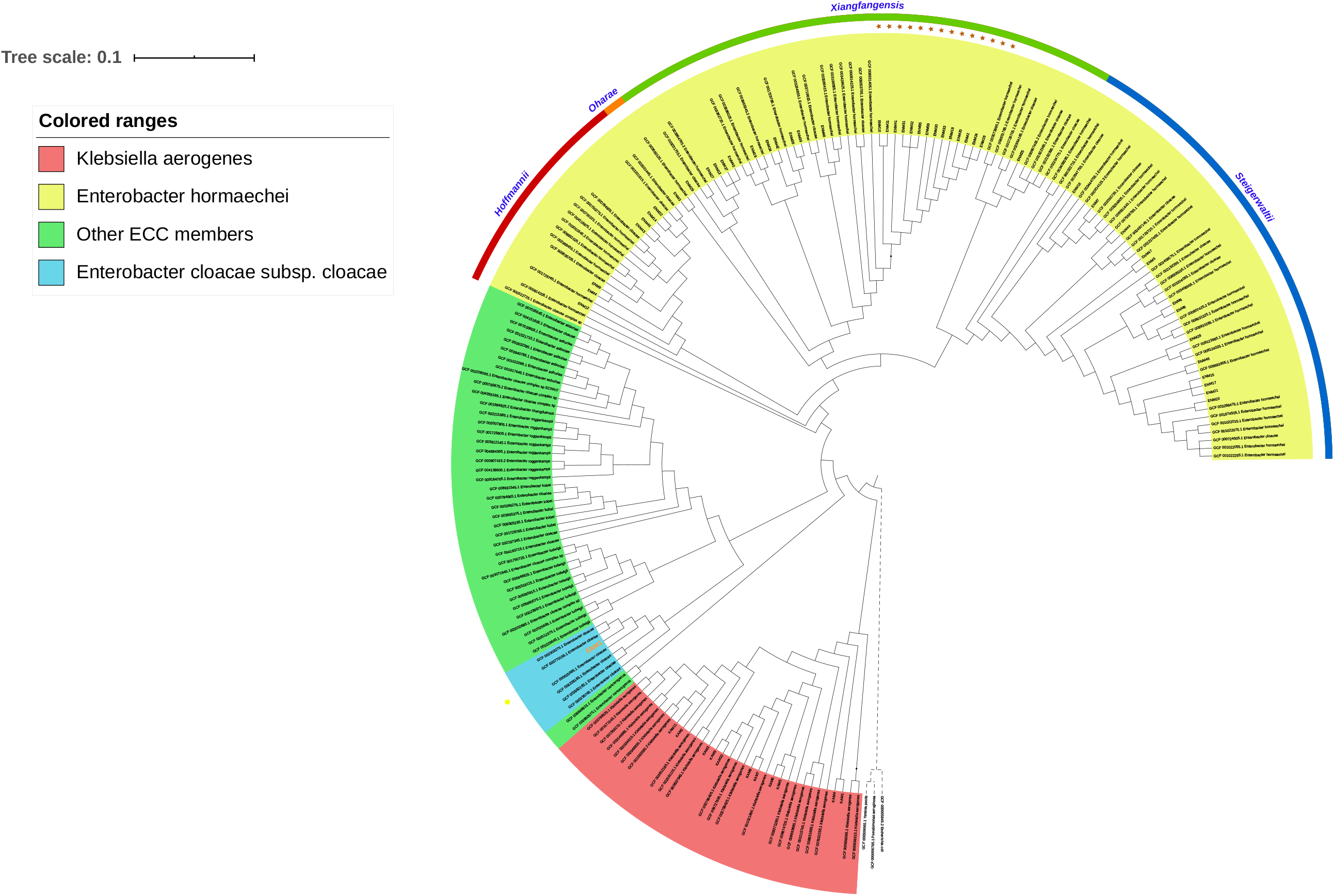
SCGs based maximum-likelihood phylogenetic tree of all study isolates and complete RefSeq ECC and *K. aerogenes* reference genomes. Colored ranges were used to differentiate between *K. aerogenes* and *Enterobacter* sp isolates. Sub-species of *E. hormaechei* are indicated by the outer rim-colored bands. Golden stars were used to highlight ST114 isolates, while the yellow dot was used to show the only *E. cloacae* subsp *cloacae* isolate (ENM1).

ECC members, except the *E. hormaechei* species, clustered in a sub-clade within clade II (green block) (Figure 5). ENM1 (blue block) was the only isolate identified as closely related to the type strain *E. cloacae* subsp. *cloacae* ATCC13047 (CP001918.1). The remaining of ECC isolates were all grouped within one sub-clade (Figure 5). Four *E. hormaechei* subspecies were detected, of which 55% (26/47) belonged to the subspecies *xiangfangensis*. Fourteen had the sequence type ST114 (Figure 5).

### Virulence of *K. aerogenes*

We looked for virulence loci in *K. aerogenes* using Kleborate, a tool to screen for ICE*Kp* (virulence-associated integrative conjugative element of *K. pneumoniae*) associated virulence loci including yersiniabactin (*ybt*), colibactin (*clb*) (Supplementary Table S3). The yersiniabactin locus (*ybtS, ybtX, ybtQ, ybtP, ybtA, ybtU, ybtT, ybtE*) encoding the yersiniabactin siderophore was detected in 36.4% (4/11) of *K. aerogenes* isolates. KAM-2, KAM-3 and KAM-11 harboured the *ybt*17 on ICE*kp10* isoform, whereas KAM-8 harboured a novel variant of the *ybt* locus. As for the bacterial toxin colibactin, all *K. aerogenes* were negative except for KAM2, KAM3, and KAM-11, which harboured the *clb3* colibactin locus.

### Plasmid identification and characterization

We used PlasmidFinder v2.1 and PCR-Based Replicon Typing (PBRT) to identify the incompatibility groups for the recovered plasmids (Figure 6). Distinct Inc groups were detected. IncFII was the most common present in 56% (n=33) of the ECC and *K. aerogenes* isolates, followed by IncFIB (33.9%; 20/59), IncX3 (22%; 13/59), IncHI1A (18.6%; 11/59), IncL (5.1%; 3/59), and IncA/C2 (1.7%; 1/59).

**Figure 6.**
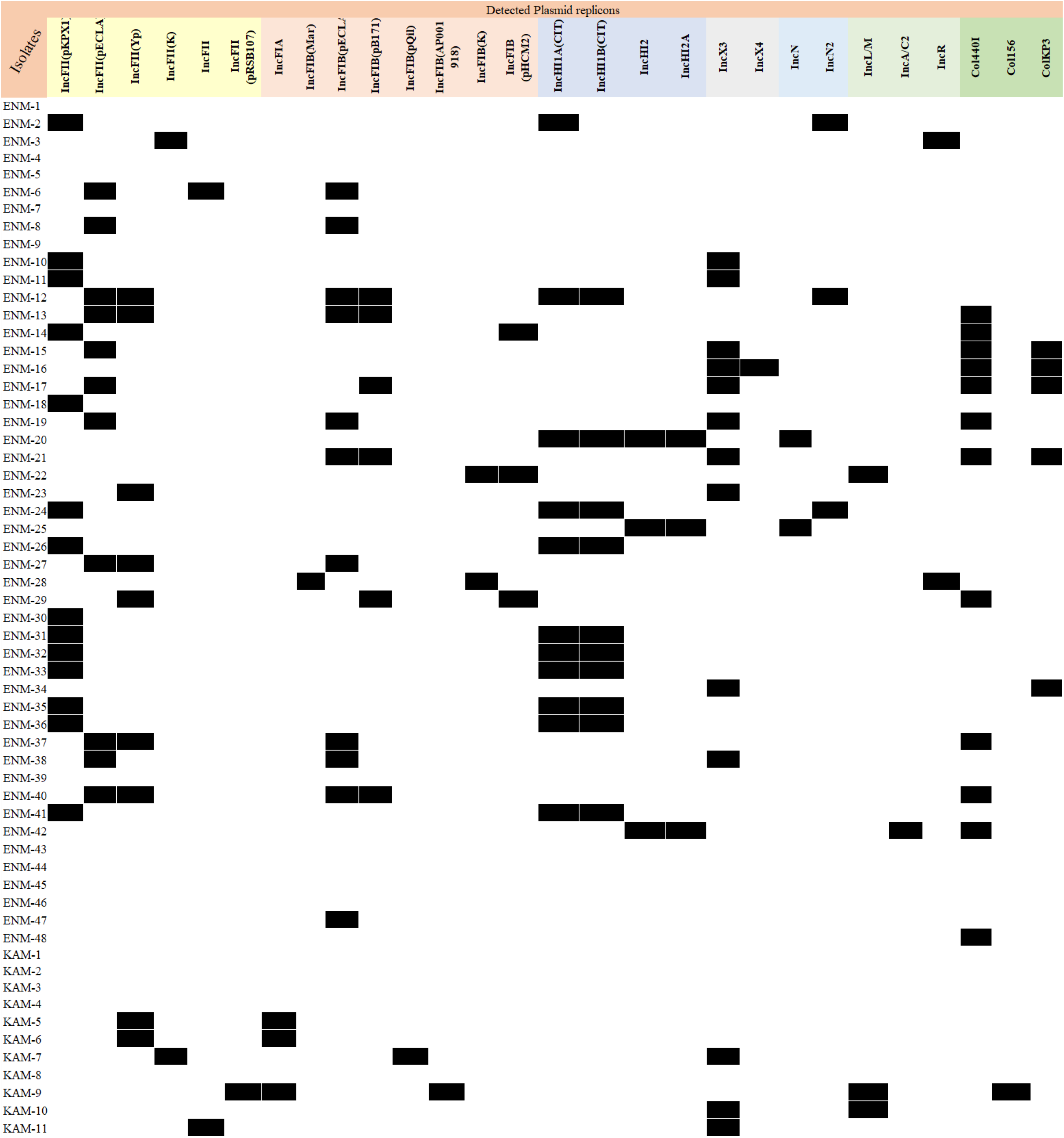
*In silico* detected plasmid incompatibility groups.

IncFII (pKPX1) was primarily associated with *bla*_NDM-1_, IncL plasmid with *bla*_OXA-48_ gene and was detected in three of the isolates (ENM-22, KAM-9, and KAM-10), and IncC which was found in a *bla*_NDM-1_ positive isolate (ENM-42) lacking all IncF groups. All *bla*_OXA-181_ harbouring isolates (ENM-15, ENM-16, ENM-17, ENM-21, and ENM-34) had the IncX3 plasmid in common and a single occurrence of *bla*_NDM-5_ was also linked to IncX3.

Three isolates, harboring the most prevalent CRDs, were selected for long read sequencing (ENM17, ENM30 and KAM9). As a result, three complete plasmids were extracted, identified and named as follows: pLAU_ENM17_OXA181, pLAU_ENM30_NDM1 and pLAU_KAM9_OXA48.

Backbone analysis of pLAU_ENM17_OXA181 (MN792919) showed that it is an IncX3 self-conjugative plasmid, harboring a *bla*_OXA-181_ class D oxacillinase, with an overall size of 51,084 bp. The backbone components of pLAU_ENM17_OXA181 were consistent with a self-conjugative plasmid, harboring the necessary genes for propagation, stability, and replication. The *bla*_OXA-181_ gene was flanked upstream by an IS*Kpn19*/transposase, followed by a replicase gene. Downstream, the class D oxacillinase was flanked by three consecutive truncated transposases ΔIS*3000*/transposase-ΔIS*Ec63*-ΔIS*26*. Moreover, the *qnrS1* fluoroquinolone resistance determinant was also identified on the plasmid and flanked upstream and downstream by IS*26* and Tn*3*-like resolvase, respectively. BLAST analysis showed 99% similarity between pLAU_ENM17_OXA181 and pSTIB_IncX3_OXA_181 (MG570092) isolated in Lebanon (36).

Based on the same analysis workflow, pLAU_KAM9_OXA48 was a 61,913 bp IncL conjugative plasmid. The IncL replicon identity was confirmed through PBRT and *in silico* BLAST analysis against publicly available IncL *repA* gene reference sequence in NCBI (NC_021488.1). Backbone screening of pLAU_KAM9_OXA48 identified conjugative transfer proteins (*tra* genes), *relB*/*relE* toxin-antitoxin system and stability related proteins. *bla*_OXA-48_ was directly bracketed by a *lysR* gene upstream and a ΔIS*1R* downstream. However, analysis of the overall genetic environment identified two IS*10A* upstream and downstream of the class D oxacillinase as follows: IS*10A*-*lysR*-*bla*_OXA48_-ΔIS*1R*-IS*10A*. Interestingly, this cassette is similar to Tn*1999*, but has different insertion sequences bracketing the cassette. IS*10A* was found to be 99% identical to IS*1999* differing in the position of two SNPs. BLAST analysis of pLAU_KAM9_OXA48 showed also showed close relatedness to other plasmids.

pLAU_ENM30_NDM1 was linked to an IncFII plasmid through *in silico* scaffolding. Detailed analysis showed the presence of a copy of *bla*_NDM-1_ directly bracketed upstream by a ΔIS*Aba125* and downstream by a *ble*_MBL_ also known as a bleomycin gene. Further analysis of *bla*_NDM-1_ genetic environment showed that it falls within a large genomic cassette harboring, from upstream to downstream: IS*3000*-ΔIS*Aba125*-*bla*_NDM-1_-*ble*_MBL_-//-Tn*5403*. BLAST analysis revealed high sequence similarity (>99%) between pLAU_ENM30_NDM1, pLAU_ENC1 (MN688131.1) (37), pAR_0128 (CP021720) and pKPX-1 (AP012055). Despite the high sequence similarity however, they were different in terms of size being larger than pLAU_ENM30_NDM1 by at least 90 Kbp.

## Discussion

In this study, we performed an in-depth molecular characterization of 48 ECC and 11 *K. aerogenes*, known previously as *E. aerogenes*, clinical isolates from Lebanon. We used molecular techniques backed by *in silico* whole-genome based characterization to resolve the molecular identity of the clinical isolates and determined and compared the phylogenetic relatedness, resistomes and virulomes. The distribution of resistance determinants was not uniform, the *bla*_NDM-1_ gene was commonly detected and associated with the IncF plasmid incompatibility group, class D oxacillinases, *bla*_OXA-48_ and *bla*_OXA-181,_ were also prevalent and were associated with IncL and IncX3 plasmid types, respectively.

ECC members and *K. aerogenes* remain largely understudied and the focus often shifts towards *E. coli* and *K. pneumoniae*. Moghnieh et al., (2019) showed the antimicrobial susceptibility data gathered from 13 local hospitals in Lebanon. Data revealed the constant increase in carbapenemase and ESBL carriage among *E. coli* and *K. pneumoniae* (16). Moreover, a study in 2018 identified 27 carbapenem resistant *E. coli* clinical isolates in a Lebanese hospital, mediated by the dissemination of the class D oxacillinases *bla*_OXA-48_, *bla*_OXA-181_ and the ESBL gene CTX-M-15 coupled with outer membrane porin alterations (38). Arabaghian et al. (2019) also characterized 34 multi-drug resistant *K. pneumoniae* clinical isolates with all being at least resistant to ertapenem (12).

Our results illustrated a predominant presence of NDM-1 (41%; 24/59), especially within the ECC population and three of the isolates harbored the *bla*_OXA-48_ gene two being *K. aerogenes* (KAM9 and KAM10). Previous reports revealed that the Middle-East and the Mediterranean regions were endemic to *bla*_OXA-48_ (12, 38, 39), and which according to our results seem to be also attributed to the prevalence of *K. aerogenes. bla*_OXA-181_, a single base pair mutant analog of *bla*_OXA-48_, was also detected in 8% (5/59) of the isolates all of which were ECC members. Both NDM-1 and OXA-181 originated from the Indian sub-continent suggesting that the driving force for carbapenem resistance within ECC in Lebanon was similar to that seen in the Balkans and the South-Eastern regions of Asia (9, 39).

Our results also highlighted the extensive occurrence of *bla*_CTX-M-15_ (49% of the isolates; 29/59), and its co-existence with other CRDs. This conforms with previous reports showing that *bla*_CTX-M-15_ was one of the most prevalent ESBL enzymes within global members of the ECC (40, 41). Haenni et al, (2016) showed that having *bla*_CTX-M-15_ among high-risk *E. cloacae* clones in companion animals, namely ST114, was alarmingly consistent (42). *bla*_CTX-M-15_ was mainly linked to mobile genetic elements but was chromosomal in ENM1. Analysis of its genetic environment revealed an IS*Ecp1* upstream and a disrupted ΔTn*2* downstream of the gene. In ENM1 the cassette was integrated downstream of the aerobactin operon which could contribute to heightening the expression of virulence and resistance determinants during host infection (43– 45).

ECC members and *K. aerogenes* are both intrinsic carriers of chromosomal *ampC* genes. *ampC* genes are inducible and prone to acquire mutations leading to their constitutive expression. When coupled with outer membrane porin alteration, primarily in *K. aerogenes*, these organisms exhibited carbapenamase independent resistance (2, 4). Both KAM5 and KAM6 were devoid of any CRDs and displayed phenotypic carbapenem resistance. Analysis of *omp36* genes showed that non-synonymous mutations were introduced through a single point mutation (C→T), introducing as a result a premature stop-codon within the gene. This is in line with the tendency of *K. aerogenes* to adapt when exposed to antimicrobial agents and promote changes in porin permeability further complicating treatment regimens (2, 10).

ST114 is an epidemic clone belonging to clonal complex CC114, and was linked to *bla*_CTX-M-15_ and recently to *bla*_NDM-1_ in Palestine, Italy, Japan, France, Spain, Greece, Latvia and other countries (9, 46). In our study, all ST114 isolates were NDM-1 carbapenemase positive highlighting the dissemination pattern of the carbapenem-resistant ECC from south-eastern Asia to Midwestern United States to the Middle Eastern countries. On the other hand, ST78 MDR *E. cloacae* was globally characterized as being a dominant ESBL clone linked to *bla*_KPC_ harboring plasmids (47). In this study two ST78 (3.4%) *E. hormaechei subsp. hoffmanni* isolates harbouring either *bla*_NDM-1_ or *bla*_OXA-48_ were detected further confirming the potential spread of high-risk clones associated with different ECC sub lineages.

The yersiniabactin and colibactin siderophores were detected in 4/11 (36.4%) and 3/11 (27.3%) of *K. aerogenes*, respectively. KAM-2, KAM-3, and KAM-11 were found to carry a conjugative and integrative isoform ICE*Kp10*. This element is associated with the mobilization of the yersiniabactin locus and coharbours the colibactin (*clb3*) and yersinbactin (*ybt17)* siderophores (48). A diverse set of ICE*Kp* isoforms were detected in carbapenem-resistant *K. pneumoniae* isolated from clinical settings in Lebanon (12). None of the multidrug resistant isolates coharboured *clb* or *ybt* proposing the positive selection for *K. aerogenes* ICE*Kp* isoform enhancing invasiveness (hypervirulence) over the resistance (12, 48).

In this study, we identified three main CRDs, *bla*_NDM-1_, *bla*_OXA-48_ and *bla*_OXA-181_ on three separate plasmids. *bla*_OXA-48_ was detected in three isolates and associated with a 61,913 bp self-conjugative plasmid pLAU_KAM9_OXA48. The plasmid being >99% similar to the reference OXA-48 plasmid pOXA-48a (JN626286.1) implicated its close relevance to the one originally recovered from *K. pneumoniae* isolated in Turkey in 2001(49). Multiple studies reported *bla*_OXA-48_ dissemination locally and globally and its association with the ∼60 Kbp widespread IncL plasmid (12, 13, 38, 50, 51). However, our study revealed a cassette-linked diversion. CRD on pLAU_KAM9_OXA48 was bracketed by IS*10A*, an isoform of IS*1999*.

OXA-181, on the other hand, a *bla*_OXA-48_-like variant, was first detected in New-Delhi in 2007 and is currently considered the second most common OXA-48 like globally. The first report of *bla*_OXA-181_ in Lebanon was from an *E. coli* (36). pLAU_ENM17_OXA181 showed >99% similarity to pSTIB_IncX3_OXA181 (recovered from Lebanon; MG570092) and pOXA181 (KP400525.1) indicating that IncX3 plasmid was highly conserved despite its wide geographic spread (36, 49, 52).

Species and sub-species within the ECC are often incorrectly identified as *E. cloacae* (53, 54). In this study only one *E. cloacae* isolate (ENM1; 2%; 1/48) was detected while the remaining were identified as being *E. hormaechei* (98%; 47/48). The predominant subspecies was the *E. hormaechei* subsp. *xiangfangensis* and was in accordance with previous reports (9, 55, 56).

To our knowledge, this is the first in-depth comprehensive whole-genome-based characterization of carbapenem resistant ECC members and *K. aerogenes* in Lebanon and the region. In this study, genome-based workflow was coupled with other more traditional approaches to build a better accurate understanding of the carbapenem-resistant population within the ECC members and *K. aerogenes* recovered from clinical settings in Lebanon. Our results revealed that the predominant sequence type among all the studied isolates was the epidemic global clone ST114 (9). The prevalent ECC identified was *E. hormaechei* subsp. *xiangfangensis* and the *bla*_NDM-1_ was the dominant detected carbapenemase. Collectively the accurate identification and targeted genome sequencing could help in reducing ineffective treatment regimens, preventing outbreaks, and limiting the spread, mobilization, and transmission of resistance determinants.

## Acknowledgments

The study was supported by the research project grants NU20J-05-00033 provided by Czech Health Research Council, by the Charles University Research Fund PROGRES (project number Q39) and by the project No. CZ.02.1.01/0.0/0.0/16_019/0000787 “Fighting Infectious Diseases” provided by the Ministry of Education Youth and Sports of the Czech Republic

All authors declare that they have no conflict of interest.

